# Peripherally administered TNF inhibitor is not protective against α-synuclein-induced dopaminergic neuronal death in rats

**DOI:** 10.1101/2024.10.11.617361

**Authors:** Josefine R. Christiansen, Sara A. Ferreira, David E. Szymkowski, Johan Jakobsson, Malú Gámez Tansey, Marina Romero-Ramos

## Abstract

The underlying cause of neuronal loss in Parkinson’s disease (PD) remains unknown, but evidence implicates neuroinflammation in PD pathobiology. The pro-inflammatory cytokine soluble tumor necrosis factor (TNF) seems to play an important role and thus has been proposed as a therapeutic target for modulation of the neuroinflammatory processes in PD. In this regard, dominant-negative TNF (DN-TNF) agents are promising antagonists that selectively inhibit soluble TNF signaling, while preserving the beneficial effects of transmembrane TNF. Previous studies have tested the protective potential of DN-TNF-based therapy in toxin-based PD models. Here we test for the first time the protective potential of a DN-TNF therapeutic against α-synuclein-driven neurodegeneration in the viral vector-based PD rat model. To do so, we administered the DN-TNF agent XPro1595 subcutaneously for a period of 12 weeks. In contrast to previous studies using different PD models, neuroprotection was not achieved by systemic XPro1595 treatment. α-synuclein-induced loss of nigrostriatal neurons, accumulation of pathological inclusions and microgliosis was detected in both XPro1595- and saline-treated animals. XPro1595 treatment increased the percentage of the hypertrophic/ameboid Iba1+ cells in SN and reduced the striatal MHCII+ microglia in the striatum of α-synuclein-overexpressing animals. However, the treatment did not prevent the MHCII upregulation seen in the SN of the model, nor the increase of CD68+ phagocytic cells. Therefore, despite an apparently positive immune effect, this did not suffice to protect against viral vector-derived α-synuclein-induced neurotoxicity. Further studies are warranted to better elucidate the therapeutic potential of soluble TNF inhibitors in PD.

## INTRODUCTION

Parkinson’s disease (PD) is a neurodegenerative disorder primarily characterized by progressive neuronal loss in various regions of the nervous system, particularly the dopaminergic neurons in the substantia nigra pars compacta (SNpc), as well as the presence of insoluble cytoplasmic alpha-synuclein (α-syn) inclusions in neurons (Forno, 1996). Additionally, PD patients exhibit microgliosis (Gerhard et al., 2006; Imamura et al., 2003; Lavisse et al., 2021) and an upregulation of pro-inflammatory markers such as IL-1β and TNF (De Lella Ezcurra et al., 2010; Dzamko, 2023; Mogi et al., 1994; Williams-Gray et al., 2016). Extensive research over the past decades indicates a pivotal role for the immune response in PD, highlighting the neurotoxic effects of a chronic pro-inflammatory environment (Tansey et al., 2022). Accordingly, epidemiological studies suggest that drugs with anti-inflammatory effects reduce the risk of PD (Alrouji et al., 2023), and genetic studies further implicate the immune system in the pathogenesis of the disease (Dzamko et al., 2015).

Interestingly, α-syn appears to be central to such immune response, since not only α-syn-induced neuronal dysfunction can ignite an immune response, but also α-syn *per se* can interact with immune receptors on innate immune cells and initiate a cellular cascade that results in the release of pro-inflammatory cytokines such as IL-1β, IL-6 and TNF (Tansey and Romero-Ramos, 2019). Accordingly, α-syn-associated pathology *in vivo* has been linked to an early neuroinflammatory response characterized by an increased number of microglia, increases in CD68 and MHCII expression, and higher levels of TNF in the nigrostriatal system (Chung et al., 2009; Sanchez-Guajardo et al., 2010; Theodore et al., 2008). Increased levels of TNF have been detected in the serum and CSF of PD patients (Eidson et al., 2017), and the excess of TNF has been extensively shown to be neurotoxic *in vivo* and *in vitro* (McCoy et al., 2011). Notably, TNF and α-syn fibrils have been suggested to compete to activate cultured astrocytes derived from induced pluripotent stem cells (iPSCs), driving distinct inflammatory transcriptomic mechanisms (Russ et al., 2021). Thus, TNF seems to be a relevant player during the α-syn-mediated neurodegenerative process in PD and a possible therapeutic target.

TNF exists in two biologically active forms: transmembrane (tmTNF) and soluble (solTNF). It is synthesized as a monomeric protein and inserted into the membrane as a homotrimer, which may then be cleaved by the TNF alpha converting enzyme (TACE/ADAM17) to form a soluble homotrimer (McCoy and Tansey, 2008). Both ligands can bind to the two homotrimeric TNF receptors: TNF receptor 1 (TNFR1) and 2 (TNFR2). The two receptors, however, differ in their expression profiles and ligand affinities; solTNF, the primary ligand for TNFR1, is constitutively expressed in most tissues; but tmTNF, expressed mainly by immune and endothelial cells, primarily signals through TNFR2 in a juxtracrine fashion (Grell et al., 1995; Grell et al., 1998). Dominant-negative TNF (DN-TNF) inhibitors are protein therapeutics or biologics consisting of modified human TNF molecules generated by introduction of discrete amino acid substitutions that disrupt receptor binding while preserving trimerization capability. Consequently, these DN-TNF variants cannot bind to either of the TNFRs but still recognize other TNF variants as self and can hetero-trimerize with them. These agents selectively inhibit solTNF via a dominant-negative mechanism by sequestering native TNF monomers and forming biologically inactive heterotrimers. In this way, a 10-fold excess of DN-TNF variant to native TNF results in a loss of >99% of native homotrimers due to monomer exchange with the mutated variants (Steed et al., 2003). Thus, the DN-TNF variants prevent solTNF-mediated intracellular signaling. The anti-inflammatory capacity and therapeutic relevance of DN-TNF biologics have been shown in models of disease with peripheral inflammatory components such as arthritis (Steed et al., 2003; Zalevsky et al., 2007), allergic airway inflammation (Maillet et al., 2011) and acute liver inflammation (Olleros et al., 2009). We have shown its neuroprotective potential in the 6-OHDA model both with intracerebral administration, but more importantly also when administered peripherally both in rat PD models and in Alzheimer’s mice models (Barnum et al., 2014; Harms et al., 2011; McAlpine et al., 2009; McCoy et al., 2006; McCoy et al., 2008).

The aim of this study was to evaluate the therapeutic potential of the peripherally administered DN-TNF therapeutic XPro1595, which selectively antagonizes solTNF, in a rat model of nigral degeneration based on viral vector-mediated overexpression of α-syn in nigrostriatal neurons. This model, unlike previous models tested with the compound, recapitulates important aspects of human PD pathology including not only the progressive loss of dopaminergic neurons but importantly the presence of α-syn pathology and neuroinflammation.

## MATERIALS AND METHODS

### Animals

Adult female Sprague-Dawley rats (*n*=35; Janvier Labs, France) weighing 225-250g upon arrival were housed in pairs in a climate-controlled facility with *ad libitum* access to food and water and were allowed to acclimate to their new housing for one week prior to starting the experiment. Throughout the study, the rats were subjected to a 12h/12h light/dark cycle; all handling of the animals was carried out during the lights-on period. The animals were weighed at least once per week throughout the study. All experimental work was performed in accordance with the Research Animal Committee at the Faculty of Health, Aarhus University and the Directive 2010/63/EU.

### Stereotaxic surgery and XPro1595 treatment

Rats received unilateral injections of recombinant adeno-associated viral vector (rAAV6) encoding either human α-synuclein (rAAV-ASYN) or green fluorescent protein (rAAV-GFP) as a control construct under control of the neuron-specific synapsin 1 promoter and the WPRE enhancer(Pircs et al., 2018). The rats were anaesthetized by i.p. injection of medetomidine hydrochloride (0.5 mg/kg) and fentanyl (0.3 mg/kg) and then placed in a stereotaxic frame (Stoelting, Wood Dale, IL, USA). An incision was made on the top of the head, and a 5 μl Hamilton syringe fitted with a glass capillary was used for injection of rAAV-ASYN or rAAV-GFP into the SN through a burr hole in the skull coordinates: 5.2 mm AP, 2.0 mm ML from bregma, and 7.2 mm DV from dura. A total of 2 μl of the viral vector solution (1×10^14^ TU/ml) was injected at a rate of 0.2 μl/30 sec. After surgery, the animals received s.c. injections of buprenorphine (0.36 mg/kg) and the antagonist atipamezole hydrochloride (0.6 mg/kg in sterile saline) before returning to their cages when fully awake. Correct placement of the needle was confirmed by immunohistochemical staining of nigrostriatal brain sections with antibodies recognizing the transgene products.

Rats received subcutaneous injections of XPro1595 provided by Xencor Inc. (Monrovia, CA, USA) at a dose of 10 mg/kg (in sterile saline) or saline as control every third day for 12 weeks starting 3 days post rAAV-surgery. The dose and dosing regimen was based on our previous study of XPro1595 in rats (Barnum et al., 2014), and other murine studies of CNS disease (Brambilla et al., 2011; Clausen et al., 2014; Novrup et al., 2014; Taoufik et al., 2011).

### Behavioral testing

*Cylinder test*: Forelimb asymmetry was evaluated using the cylinder test at 6 and 12 weeks post-surgery, as previously described (Tentillier et al., 2016). Briefly, the rat was placed inside a glass cylinder (20 cm in diameter, 30 cm height) and was allowed to move freely; during exploration of the cylinder, the rat would spontaneously stand on its hind legs and touch the cylinder wall with the forepaws. The test was videotaped, and an observer blind to the animal’s group/treatment counted the number of contacts made by each forelimb with the cylinder wall. In total, 20 contacts were counted per animal. A contact was defined as touching of the cylinder wall with the entire palm. Data are presented as the number of contacts with the contralateral forelimb as a percentage of total.

*Stepping test*: The stepping test, which is used for assessment of forelimb hypokinesia, was performed 11 weeks after the surgery. Briefly, the experimenter held the rat with one forepaw fixed and the other free to move. The rat was then moved along a horizontal surface (80 cm) for approximately 8 sec during which the rat made adjusting steps with the free-moving paw on the surface. This was repeated four times with each paw in both the forehand (i.e., the animal is moved left when the right paw touches the table and vice versa) and backhand direction (i.e., the animal is moved right when the right paw touches the table and vice versa). The rats were trained for two days prior to the actual test. The test was videotaped, and an observed blind to the animal’s identity counted the number of adjusting steps made by the rat as it was moved along the surface. Stepping test data are presented as the number of steps using the contralateral paw as percentage of the number of steps using the ipsilateral paw in the backhand direction.

### Perfusion-fixation and tissue processing

Twelve weeks after rAAV-surgery, the rats were deeply anaesthetized with an overdose of pentobarbital (250 mg/kg). Upon respiratory arrest, the thorax was opened and cardiac blood collected. The spleen was collected, immediately snap frozen on dry ice, and stored at -20**°**C. Subsequently, the animal was intracardially perfused first with ice-cold 0.9% saline, followed by 4% paraformaldehyde (PFA) in 0.1 M sodium phosphate buffer. The brain was removed and post-fixed in 4% PFA for 3 hours at 4**°**C and thereafter immersed in 25% sucrose (with 0.05% sodium azide) at 4**°**C until sectioning into 40 μm-thick coronal sections on a freezing microtome (HM 450 Sliding Microtome, Microm Int. Gmbh, Walldorf, Germany). Tissue sections were stored in anti-freeze solution (30% ethylene glycol, 30% glycerol) at -20**°**C until histological processing.

### Immunohistochemistry

Immunohistochemical staining was performed on free-floating brain sections as before (Tentillier et al., 2016). The sections were then incubated with primary antibodies for: human α-synuclein (Hu-α-syn, 1:4000, Abcam #ab138501), GFP (1:30000, Abcam #ab290-50), anti-phosphorylated α-synuclein (pSer129 α-syn, 1:3000, Cell Signaling Technologies #23706), vesicular monoamine transporter-2 (VMAT-2, 1:5000, Everest Biotec #EB06558), Tyrosine Hydroxylase (TH, 1:3000, Merck Millipore #MAB318), MHCII (1:250, BioRad #MCA46G), Iba1 (1:800, Wako Fujifilm #019-1974), CD68 (1:200, AbD Serotec #MCA341) and GFAP (1:5000, Abcam #ab7260) in 2.5% serum in T-KPBS overnight at room temperature. Next, brief blocking was performed with 1% serum in T-KPBS for 10 minutes at room temperature, followed by incubation with an appropriate biotinylated secondary antibody (goat anti-rabbit, horse anti-goat, or horse anti-mouse; Vector Laboratories, Burlingame, CA, USA) diluted 1:200 in T-KPBS for two hours at room temperature. Subsequently, the sections were incubated with avidin-biotin-peroxidase complex (ABC Elite; Vector Laboratories, Burlingame, CA, USA) in KPBS for one hour at room temperature. Finally, immunostaining was developed with 0.05% 3,3’-diaminobenzidine (DAB) and 0.1-1% H_2_O_2_. The sections were mounted on chromium-potassium-gelatine-coated slides, dehydrated in increasing concentrations of ethanol, cleared in xylene, and coverslipped with DPX mountant (06522; Sigma-Aldrich, St. Louis, MO, USA). Cresyl violet was used for nuclear counterstaining of sections immunostained with anti-MHCII antibody.

### Stereological quantification of cells

Stereological quantification of VMAT2+ neurons and Iba1+ cells in SN were performed by an observer blind to the animal’s identity, using the optical fractionator principle on a Leica DM6000 B microscope. Counting was performed in every 6^th^ section of the SN (8-10 sections per animal, spaced 240 μm). A 1.25x objective lens was used to outline the borders of the SN, and counting was performed using a 63x oil objective (NA 1.25) and VIS newCAST software (Visiopharm Integrator System v5.0.5.1399, Visiopharm A/S, Hørsholm, Denmark). The counting frame (56.89μm x 42.66 μm) was randomly located by the VIS module and methodically moved to sample the entire delineated region of the SN, with a step length of 125-200 μm yielding an average final count of 150 cells on each side of the brain. The total number of VMAT2+ cells was estimated using the optical fractionator formula (Gundersen and Jensen, 1987; West, 1999). A coefficient error (CE) <0.10 was accepted.

Iba1+ ramified microglia were counted with a 40X objective, and a step length of 225-275 μm, which assured that at least 200 cells were counted on each side of the brain in each animal. Four Iba1+ cellular profiles were defined as previously described [8]. The percentage of each morphological cell type was calculated as the total % of A, B, C, and D type in each section and averaged per animal. The estimated number of Iba1+ cells was calculated according to the optical fractionator formula.

### Measurement of optical fiber density

Striatal TH-or VMAT2-immunoreactive fiber density was determined by optical densitometry. Immunostained sections were scanned in grayscale using an Epson Perfection 3200 Photo scanner, and optical density was measured blindly using ImageJ 1.49v software (National Institutes of Health, USA). For each section, the optical density of the striatum was normalized to the density of corpus callosum since this structure is devoid of any TH or VMAT2 immunosignal. To determine TH+ fiber density, 6 sections separated by 480 μm were analyzed corresponding approximately to +1.20, +0.70, +0.20, -0.30, -0.80, -1.30 mm relative to bregma according to the rat brain atlas by Paxinos and Watson (Paxinos and Watson, 2007). For VMAT2+ fiber density, 4 sections spaced 960 μm were analyzed corresponding approximately to +2.00, +1.00, 0.00, -1.00 mm relative to bregma.

### Quantification of striatal TH+ fiber swellings

For each animal, the number of TH+ fiber swellings in the striatum was determined in three striatal sections. In total, six photos were taken of the ipsilateral side at the lateral (1L, 2L), or medial striatum (1M, 2M) for the two most frontal sections, and one dorsal (3D), and another ventral (3V) for the most caudal section. TH+ swellings ≥2 μm^2^ were quantified using ImageJ 1.49v software with a standard brightness threshold of 115 and circularity 0.30-1. The data are presented as the total number of swellings (≥2 μm^2^) as well as the number of small (2-5 μm^2^), medium-sized (5-10 μm^2^), and large swellings (>10 μm^2^) per area.

### Quantitative analysis of α-syn phosphorylation, MHCII+/CD68+ microglia and GFAP+ astrocytes

In order to evaluate the α-syn pathological phosphorylation and the associated microglial and astrocytic response, the expression of pSer129 α-syn, MHCII+/CD68+ microglia and GFAP+ astroglia in the nigrostriatal pathway were assessed. *MHCII*: Within the striatum, all ramified MHCII+ cells were counted in every 12^th^ section (7 sections per animal). In addition, the magnitude of microglial activation in the SN was assessed through quantification of the area covered (%Area) by MHCII+ immunostaining using the Upright Widefield Slide Scanner Microscope (UWSSM, Olympus VS120) at 20× magnification. Three Enhanced Focal Images (EFI) with 20 μm depth (z-axis) were captured from three equally distant coronal sections located between -4.80 and -6.04 mm from bregma. EFI photos were analyzed with ImageJ (Fiji) software using a protocol for unbiased automated counting of ramified MHCII+ cells. *pSer129*, *CD68* and *GFAP*: Three EFI pictures were captured as described above in three striatal sections (between 1.60 and -0.80 mm from bregma) and three nigral sections (between -4.80 and -6.04 mm from bregma) and analyzed with ImageJ (Fiji) software. The same settings were applied to all animals from each immunostaining. The %Area occupied by the positive staining was averaged and compared across groups.

### Statistical analysis

Statistical analysis was performed using Prism 6.0 (GraphPad Software Inc., San Diego, CA, USA). Gaussian Normality was first assessed using normality distribution with a Shapiro-Wilk normality test followed by a two-way analysis of variance (ANOVA) and, when appropriate, by a post-hoc Tukey’s multiple comparisons test. Outliers were addressed and excluded by the ROUT method (Q=1%). All histograms are presented as mean ± standard deviation (SD). In all cases, a test result was considered significant when p<0.05.

## RESULTS

To investigate the neuroprotective potential of peripheral injections of XPro1595 towards α-syn-driven dopaminergic neurodegeneration, we injected rAAV α-syn, or rAAV GFP control, in the rat midbrain (serotype 6, synapsin 1 promoter and the WPRE enhancer), and three days post-injection administered XPro1595 s.c. every third day for the following 12 weeks (**FIG. 1A**). No obvious adverse side effects were found, and all groups showed similar weight gain trough the experiment **(FIG. S1A**). We examined macroscopically the spleens to evaluate possible affection of secondary lymphoid organs and verified that XPro1595 treatment did not affect the spleen size or appearance **(FIG. S1B)**.

**FIG. 1.**
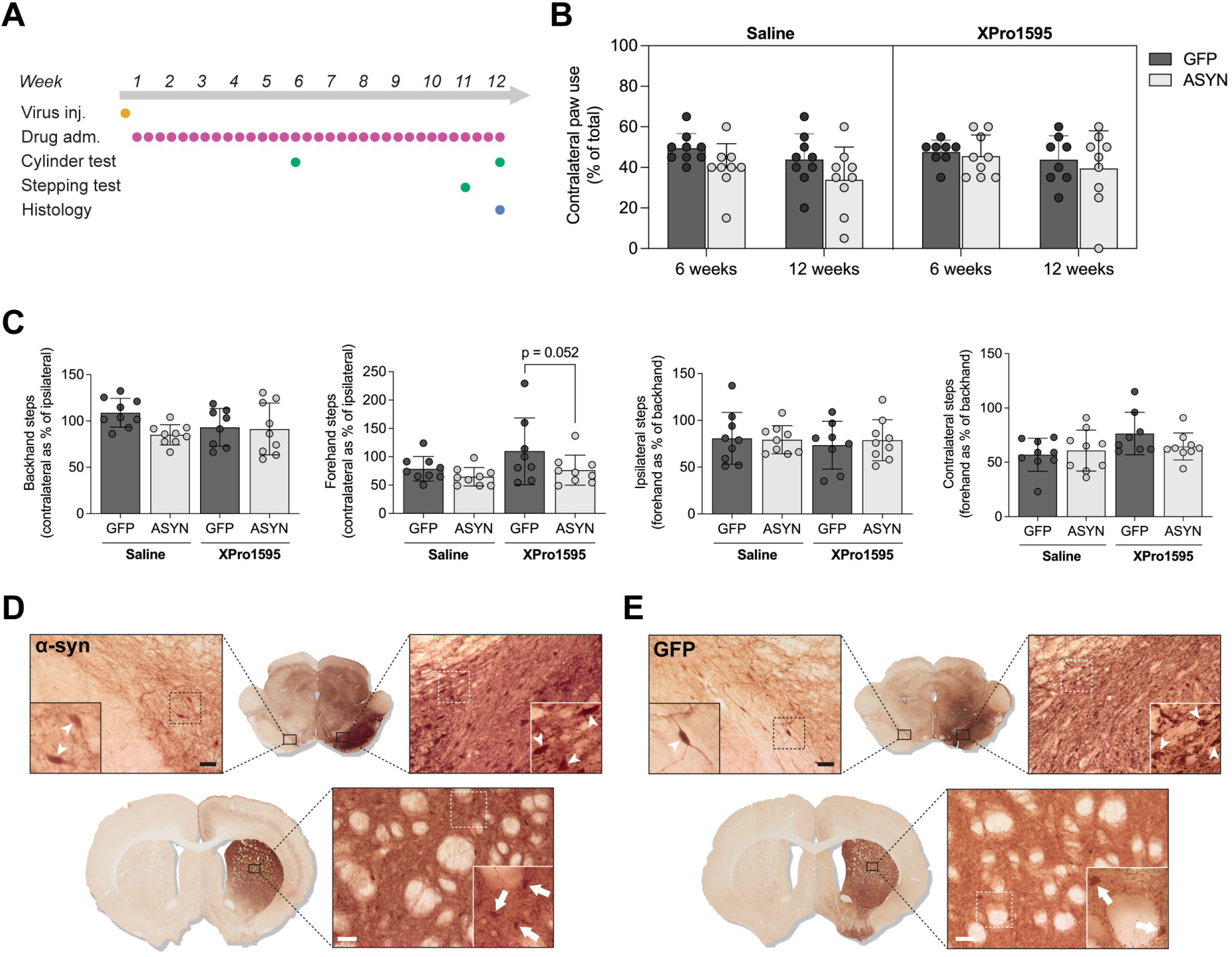
Transgene expression and motor performance. Assessment of the functional outcome of the XPro1595 treatment was done using two different behavioral tests designed to detect deficits in motor performance. **A** Outline of the study. **B** The cylinder test was performed at 6 weeks and 12 weeks post-rAAV surgery. **C** The stepping test was performed at 11 weeks post-surgery. Histograms present the mean ± SD; two-way ANOVA followed by Tukey’s multiple comparisons test (*n* = 8-9/group). **D-E** Representative images of α-syn (**D**) and GFP (**E**) immunostaining of the substantia nigra (SN) (upper panels) and striatum (lower panels). Arrowheads indicate robust transgene expression in nigral neurons in the ipsilateral side with limited dissemination to the contralateral side. Projections in the ipsilateral striatum showed strong immunosignal, and the presence of immunoreactive striatal cell bodies (arrows) indicates anterograde spreading of the rAAV vector. Scale bar: 50 µm.

### α-syn overexpression did not induce robust motor impairment

Analysis of motor behavior was done using the cylinder test (6 & 12 weeks post-surgery) and the stepping test (11 weeks). The cylinder test showed no significant bias in the use of the forelimb at any time point **(FIG. 1B)**. In the stepping test, α-syn overexpression resulted in a clear trend towards a reduced number of backhand steps in the ASYN/Saline rats (p=0.069) when compared to the GFP/Saline animals **(FIG. 1C)**. This effect was not observed in the XPro1595 group, although there was a significant difference between GFP and ASYN in the forehand steps. This appeared to be related to a higher number of steps in the GFP/XPro1595 group rather than to a treatment effect on the ASYN group **(FIG.1C)**. Thus, α-syn overexpression resulted in very mild motor changes, that were not improved by the XPro 1595.

### Systemic solTNF inhibition did not prevent axonal pathology induced by α-syn overexpression

Twelve weeks post-rAAV injections all animals showed robust expression of α-syn or GFP, respectively, in neurons in the ipsilateral SN with some expression in the contralateral side **(FIG. 1D-E**, **arrowheads).** Specific immunostaining of striatal fibers proved efficient transport of the transgene protein along the nigrostriatal axons **(FIG. 1D-E**, **lower panels)**. Notably, striatal cell bodies expressing α-syn or GFP, respectively, were observed in all animals indicating anterograde spreading of the viral particles **(FIG. 1D-E**, **arrows)**. Likewise, at the level of the midbrain, transgene expression was not confined exclusively to the SN, indicating some degree of diffusion of the viral vector from the injection site to adjacent areas in the ipsilateral side **(FIG.1D-E, upper panels)**. Microscopy analysis of striatal fibers in the ipsilateral striatum of ASYN animals revealed a pathological appearance with dystrophic fibers presenting α-syn+ swellings and fiber thickening. These dystrophic fibers contained phosphorylated α-syn as shown by immunostaining with an anti Ser129 phospho-α-syn **(FIG.2A, C)**, which also found cell bodies in the ipsilateral SN of the ASYN-overexpressing animals **(FIG.2B, D)** which was not prevented by XPro1595 treatment. Such axonal changes were not observed in GFP overexpressing animals, or in the contralateral side of any group **(FIG.2A-D)**.

The pathological axonal terminal swellings induced α-syn overexpression were also positive for TH and VMAT2 **(FIG. 2E, white arrowheads)**, irrespective of the treatment but it was absent in all GFP animals. To evaluate the degree of axonal damage, we quantified the number of small (2-5 µm^2^), medium-sized (5-10 µm^2^), and large (>10 µm^2^) round TH+ swellings at two medio-lateral areas in three different rostro-caudal sections **(FIG. 2D-F)**. In both ASYN groups, TH+ swellings were generally more pronounced in the rostral part of the striatum and gradually decreased in the caudal direction. At the two most rostral levels, the swellings were more abundant in the lateral striatum **(FIG. 2F)**. In the ASYN/XPro1595 group, a small, non-significant decrease was observed in the number of small and medium-sized swellings in the two most frontal sections compared to the ASYN/Saline group **(FIG. 2F)**, while the number of swellings in the caudal part was comparable in the two groups. This might be due to a lower axonal density observed in frontal sections in the ASYN/XPro1595 group **(FIG. 3)**.

**FIG. 2.**
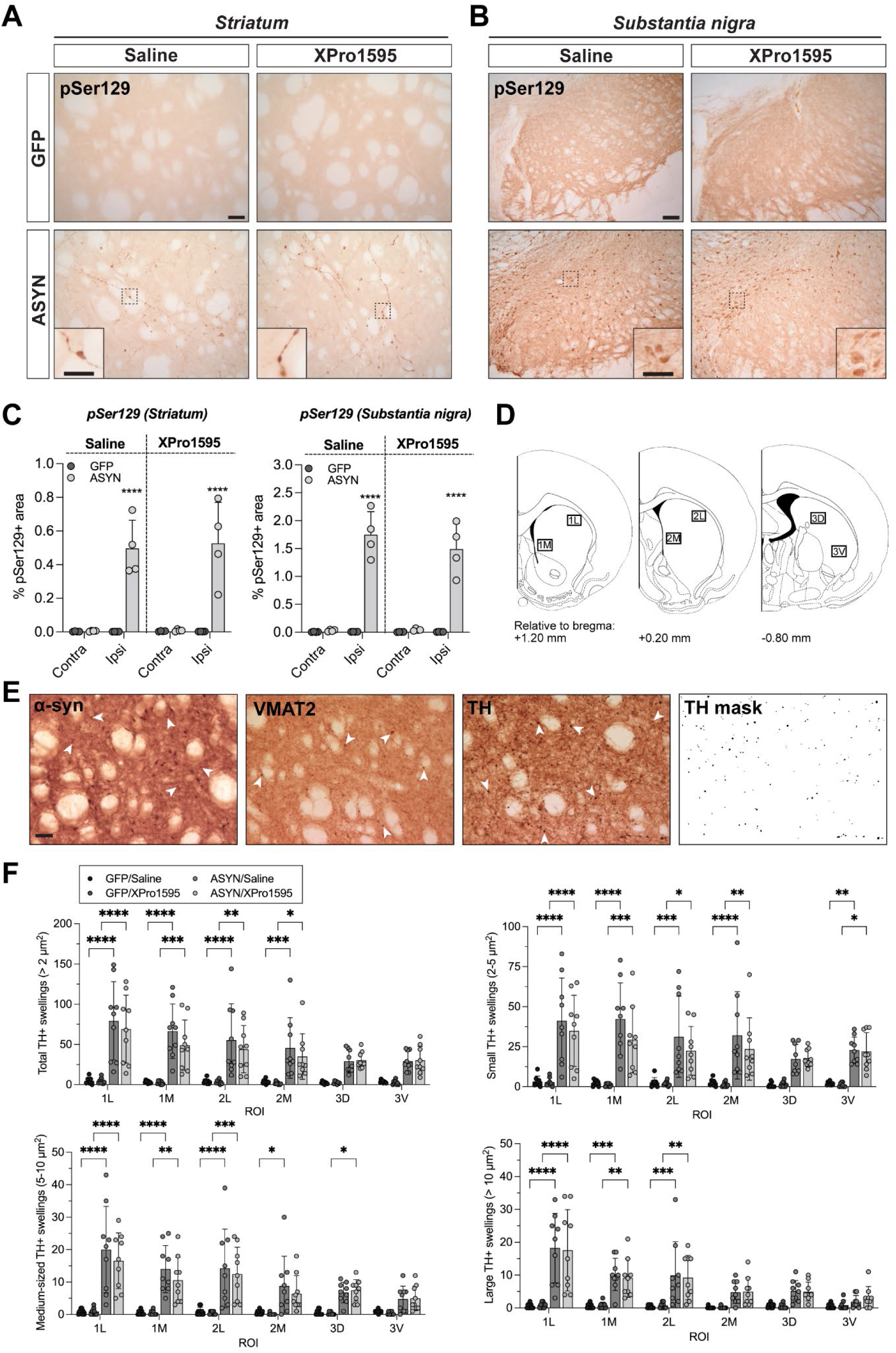
Pathological α-syn aggregation in the nigrostriatal pathway. **A-B** Representative images of pSer129 α-syn aggregates in the ipsilateral striatum and SN, respectively. Scale bar represents 50 µm for striatum (25 µm for respective cropped image) and 100 µm for SN (50 µm for respective cropped image). **C** Bar graphs with individual values illustrate the % of area covered by pSer129+ immunostaining in the contralateral and ipsilateral striatum and SN. The histograms present the mean ± SD (*n* =4/group). Two-way ANOVA followed by Sidak’s multiple comparisons test; *p<0.05, **p<0.01, ***p<0.001, ****p<0.0001. **D** The number of TH+ fiber swellings was determined in three striatal sections. In total, six photos were taken in the ipsilateral side at the lateral (1L, 2L), or medial striatum (1M, 2M) for the two most frontal sections, and one dorsal (3D), and another ventral (3V) for the most caudal section (left). **E** Example images of the dorsolateral striatum (corresponding to 1L) immunostained against α-syn (left), VMAT2 (middle), and TH (right) from a representative ASYN/Saline animal. All markers show some degree of fiber swelling (arrowheads). Right-most image shows the mask of the TH-immunostained section used for quantification of terminal swellings ≥2 µm^2^. **F** Bar graphs showing the total number of TH+ swellings (≥2 µm^2^) as well as the number of small (2-5 µm^2^), medium-sized (5-10 µm^2^), and large (>10 µm^2^) swellings in the six striatal areas. The histograms present the mean ± SD (*n* = 8-9/group). For each area, a two-way ANOVA was performed followed by Tukey’s multiple comparisons test; asterisk(s) indicate statistical difference from the respective GFP control group (*p<0.05, **p<0.01, ***p<0.001, ****p<0.0001).

**FIG. 3.**
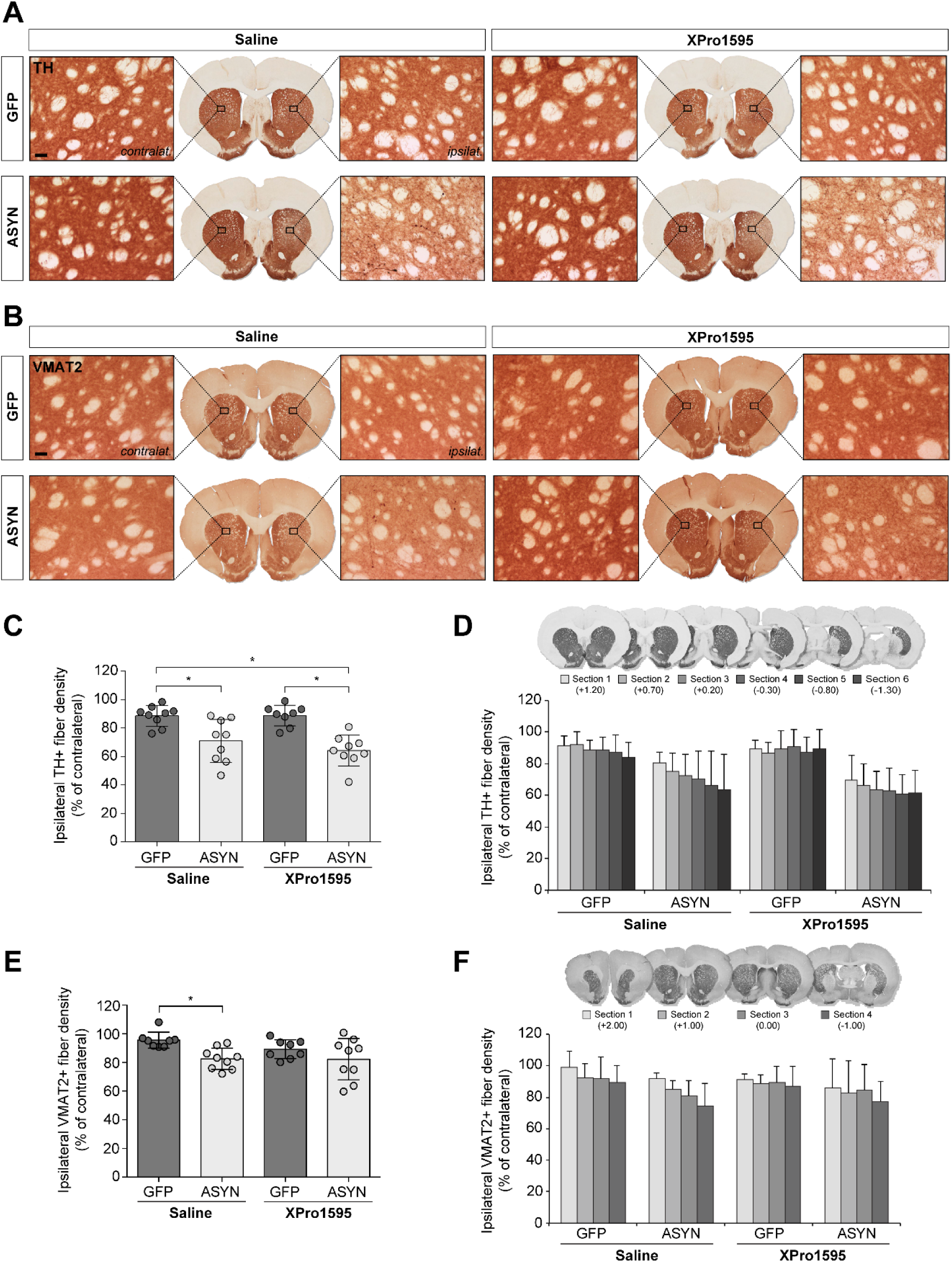
Striatal dopaminergic fiber density. **A-B** Representative images from each experimental group showing striatal TH+ (**A**) and VMAT2+ (**B**) neuronal terminals in the contralateral (left) and ipsilateral side (right). Scale bar: 50 µm (applies to all). **C** Bar graph showing the ipsilateral TH+ fiber density as percentage of the contralateral side in animals receiving rAAV-GFP (dark gray) or rAAV-ASYN (light gray). **D** Representative images of six TH immunostained striatal sections used for determination of dopaminergic fiber density. Bar graphs showing the ipsilateral fiber density as percentage of the contralateral side in each section across the rostro-caudal axis (increasingly dark nuances in the caudal direction). The histograms present the mean ± SD (*n* =8-9/group). Two-way ANOVA followed by Tukey’s multiple comparisons test; *p<0.05, ****p<0.0001. **E** Bar graph showing the ipsilateral VMAT2+ fiber density as percentage of the contralateral side in animals receiving rAAV-GFP (dark gray) or rAAV-ASYN (light gray). **F** Representative images of four VMAT2 immunostained striatal sections used for determination of dopaminergic fiber density. Bar graphs showing the ipsilateral fiber density as percentage of the contralateral side in each section across the rostro-caudal axis (increasingly dark nuances in the caudal direction). The histograms present the mean ± SD (*n* =8-9/group). Two-way ANOVA followed by Tukey’s multiple comparisons test; *p<0.05.

### Decrease in dopaminergic fiber density was not prevented by solTNF inhibition

To evaluate the integrity of nigrostriatal projections and the possible treatment effect, densitometric analysis of the striatal dopaminergic innervation was performed using two markers TH **(FIG. 3A)** and VMAT2 **(FIG. 3B)**. α-Syn overexpression in saline-treated animals led to an ipsilateral reduction of the striatal TH+ fiber density (29.0±15.1% vs. contra), which was significantly greater than that observed in the respective GFP control (11.4±7.2%) **(FIG. 3A, C)**. This decrease in TH+ fiber density was not prevented by the DN-TNF treatment as ASYN/XPro1595 rats also showed a significant TH+ fiber loss (35.8±10.7%) when compared to the GFP/XPro1595 control (11.2±7.4%).

VMAT2+ fiber density confirmed the α-syn-induced reduction of dopaminergic fibers in the ipsilateral striatum, although of a lower magnitude than for TH+ fibers **(FIG. 3B, E)**. The saline- and XPro1595-treated ASYN groups showed a comparable decrease in ipsilateral VMAT2+ fiber density (17.5±7.4% and 17.8±14.5%, respectively), which was significantly greater than the loss observed in the GFP/Saline group (6.7±8.6%). However, the ipsilateral VMAT2+ fiber density measured in the ASYN/XPro1595 group was not significantly different to that in the GFP/XPro1595 control that showed a variable level of denervation in some animals (average -10.8±6.6%).

As previously described (Phan et al., 2017) the ASYN/Saline group showed a rostro-caudal gradient in the loss of both TH+ and VMAT2+ fibers with the caudal part being mostly affected **(FIG. 3D,F)**. This was less pronounced in the ASYN/XPro1595 group that showed comparable reduction at all rostro-caudal levels. No obvious differences were observed between the two GFP control groups, which both showed a slight decrease in fiber density at all rostro-caudal levels.

### SolTNF inhibition failed to prevent the loss of dopaminergic neurons in substantia nigra

To measure whether DN-TNF treatment could protect against α-syn overexpression-induced neuronal loss we performed stereological quantification of VMAT2+ cells in SN **(FIG. 4)**. α-syn overexpression induced a significant loss of VMAT2+ neurons in the ipsilateral side in saline-treated animals compared to the contralateral side (33.0±13.1%), which was not averted by systemic treatment with XPro1595 (42.2±14.1%). A small, yet significant, ipsilateral cell loss was seen in both saline- and XPro1595-treated GFP animals (18.6±12.7% and 19.4±11.1%, respectively). However, the number of VMAT2+ neurons in the ipsilateral SN was always lower in ASYN animals compared to the ipsilateral SN of the correspondent GFP control (**FIG. 4B**). Accordingly, when expressing the number of ipsilateral VMAT2+ neurons as percentage of the contralateral side, both ASYN groups showed a greater cell loss compared to their GFP control group **(FIG. 4C)**.

**FIG. 4.**
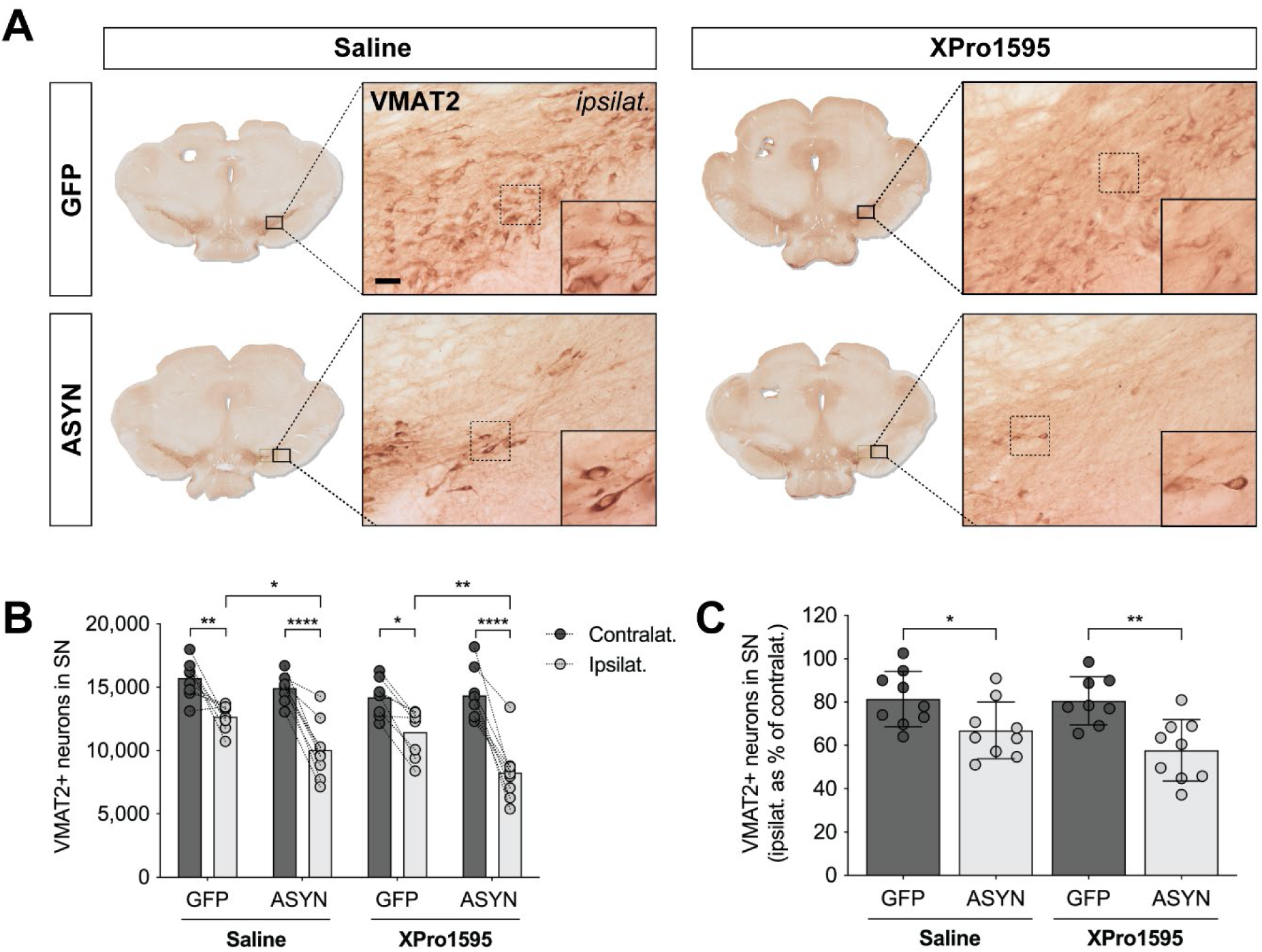
Stereological quantification of dopaminergic neurons in the Substantia nigra. **A** Representative images showing VMAT2+ neurons in SN from each experimental group. Scale bar: 50 µm (applies to all). **B** Bar graph showing the estimated number of VMAT2+ neurons in SN in the contralateral (dark gray) and ipsilateral (light gray) side. **C** Bar graph showing the number of VMAT2+ neurons in the ipsilateral side as percentage of the contralateral side in animals receiving rAAV-GFP (dark gray) or rAAV-ASYN (light gray). Scale bar: 12.5 µm. The histograms present the mean ± SD (*n* =8-9/group). Two-way ANOVA followed by Tukey’s multiple comparisons test; *p<0.05, **p<0.01, ***p<0.001, ****p<0.0001.

As previously mentioned, the decrease in TH+ striatal fiber density was greater than that seen for VMAT2+ fibers. Nevertheless, the TH+ and VMAT2+ fiber densities were positively correlated **(FIG. S2A)**. Moreover, the TH+ fiber density correlated positively with the number of VMAT2+ neurons in the SN **(FIG. S2B)**.

### SolTNF inhibition modulated the microglial response to α-syn overexpression

We investigated the possible effects of the XPro1595 treatment on the α-syn-induced neuroinflammatory reaction, by quantifying and profiling the Iba1+ microglia cells in the SN using stereology **(FIG. 5A-E)**. We have previously shown that α-syn overexpression in the rat midbrain induces an ipsilateral proliferation of microglia that peaks at 2 months to decrease later but without reaching contralaleral levels after 15 weeks (Sanchez-Guajardo et al., 2010). Here, we found that α-syn overexpression also induced significant ipsilateral increase in total Iba1+cells (+21,77% vs. contra), which was also significantly higher than the number of Iba1+ cells in the GFP/saline ipsilateral SN **(FIG. 5B)**. Systemic XPro1595 treatment did not prevent the ipsilateral microgliosis (+18,64%), which was also significant in the GFP/XPro1595 group (+16,37% vs. contra, **FIG. 5B**).

**FIG. 5.**
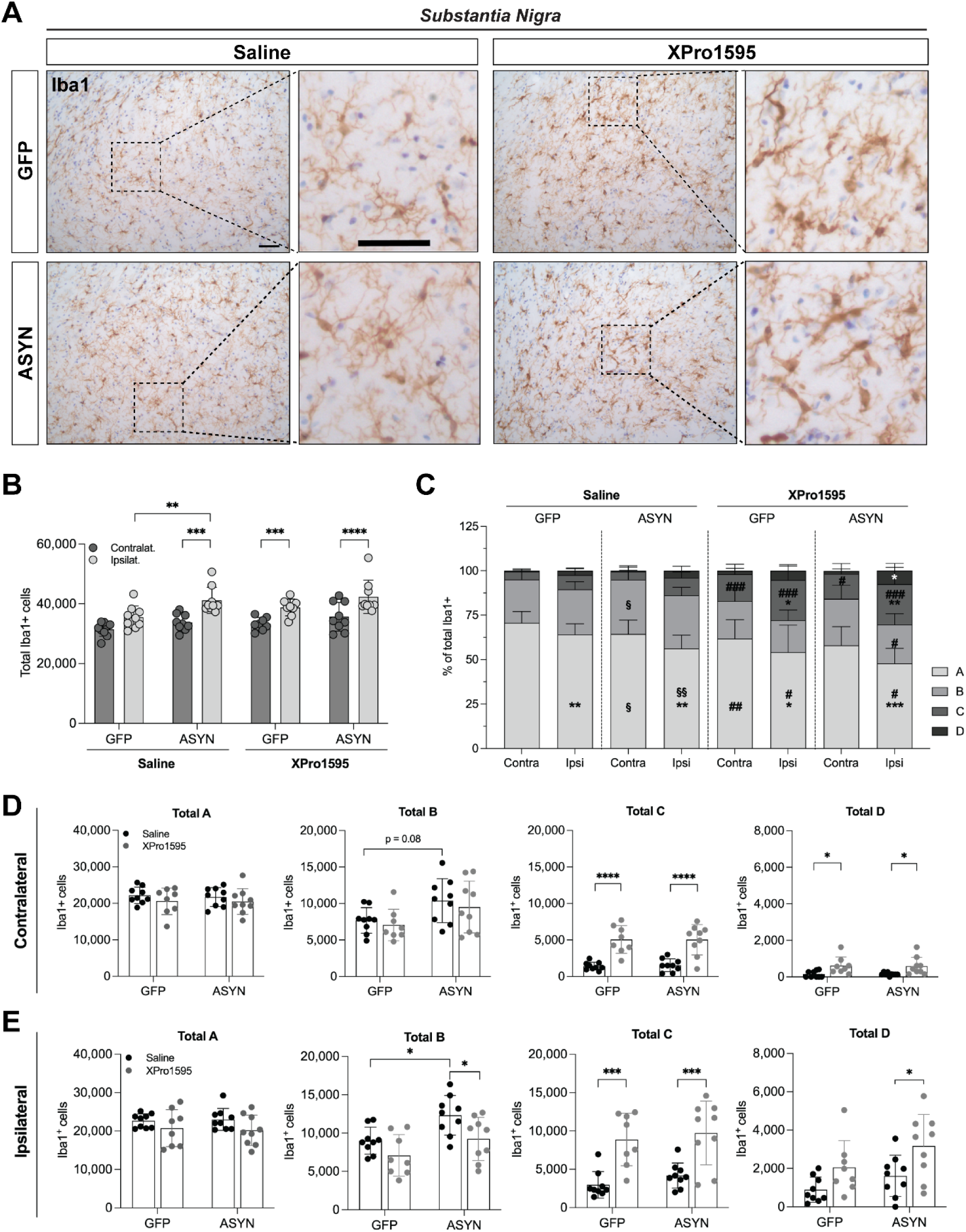
Iba1+ microglia in the Substantia nigra. **A** Representative images of Iba1+ cells in the ipsilateral SN. Scale bar represents 50 μm. **B** Bar graph represent the total number of Iba1+ cells in the contralateral and ipsilateral SN. **C** Stacked bar graph illustrates the percentage of Iba1+ cell subtypes/states (A, B, C and D) in the contralateral and ipsilateral SN. *represents difference to contralateral; # represents difference to treatment (same rAAV); § represents difference to rAAV (same treatment). Total contralateral and ipsilateral cell subtypes are represented in the bar graphs in **D** and **E**, respectively. The histograms present the mean ± SD (*n* =8-9/group). Two-way ANOVA followed by Sidak’s multiple comparisons test; *p<0.05, **p<0.01, ***p<0.001, ****p<0.0001.

We also analyzed the morphology of Iba1+ as profile changes are associated to microglia activation. In short, surveillant Type A and ramified Type B are the most common microglia profile found in healthy brain; hypertrophic Type C and ameboid Type D are very rarely seen in healthy brain, but are associated with neuroinflammatory conditions as seen previously in this model (Sanchez-Guajardo et al., 2010). Analysis of the different morphological Iba1+ cell profiles revealed a decrease in the percentage of the homeostatic type A (% of total Iba1+) in the ipsilateral SN of both GFP and ASYN saline (vs. contralateral) **(FIG. 5C-E)**, but this decrease was greater in the ASYN (vs. GFP) **(FIG. 5C)**. α-Syn overexpression (ASYN/saline) led to a significant expansion in the total number of Type B and to non-significant increase in the hypertrophic Type C and ameboid Type D in the ipsilateral side (vs.GFP/saline) **(FIG. 5C,E)**. Thus, α-syn overexpression led to more activated morphological profiles of Iba1+ cells and a decrease in the surveillant type A. Systemic XPro1595 treatment led to further decrease of the type A percentage in the ipsilateral SN (vs. saline) **(FIG. 5C)** and an increase in the number of hypertrophic type C on both SN sides irrespective of the rAAV injected **(FIG. 5C, E-Total C)**. Furthermore, XPro1595 treatment increased the ameboid type D in the ipsilateral SN only in ASYN animals (% of total & total numbers **(FIG. 5C, E-Total D)** in detriment to the type B microglia, which was decreased (vs. ASYN/saline) **(FIG. 5C, E-Total B)**. Therefore, chronic treatment with XPro1595 resulted in an increase of the hypertrophic microglia in all animals, and of the ameboid microglia type D in the ASYN overexpressing rats.

To further assess the effect of XPro1595 treatment on the immune response, we analyzed MHCII and CD68 in both striatum and SN (**FIG. 6 & 7**). α-syn overexpression led to increased MHCII expression in the SN **(FIG. 6A,B)**, whereas GFP expression led to a more modest response with only some MHCII+ cells scattered throughout the SN *pars compacta*. In the striatum, MHCII+ microglia were found either as individual cells or in small clusters of 2-4 cells **(FIG. 6D)**, mainly A- and B-type ramified MHCII+ cells with occasional C- and D-type were observed in all groups **(FIG. 6D,E)**. Occasional round MHCII+ cells in the parenchyma or associated with a blood vessel were found in all groups, suggesting infiltration of peripheral cells, potentially T-cells **(FIG. 6F).** MHCII+ microglia were consistently sparse in the contralateral side **(FIG. 6C)**. The ASYN/Saline group showed a 2-fold increase in the number of MHCII+ microglia in the ipsilateral side (relative to contralateral, **FIG. 6C**), which was significantly higher than the ipsilat./contralat. ratio observed in the GFP/Saline group. XPro1595 treatment attenuated this increase in striatal MHCII+ microglia, and the number of MHCII+ cells in the ASYN/XPro1595 was similar to that in GFP/XPro1595 animals, suggesting that the DN-TNF may have prevented the α-syn-induced upregulation of MHCII in striatal microglia.

**FIG. 6.**
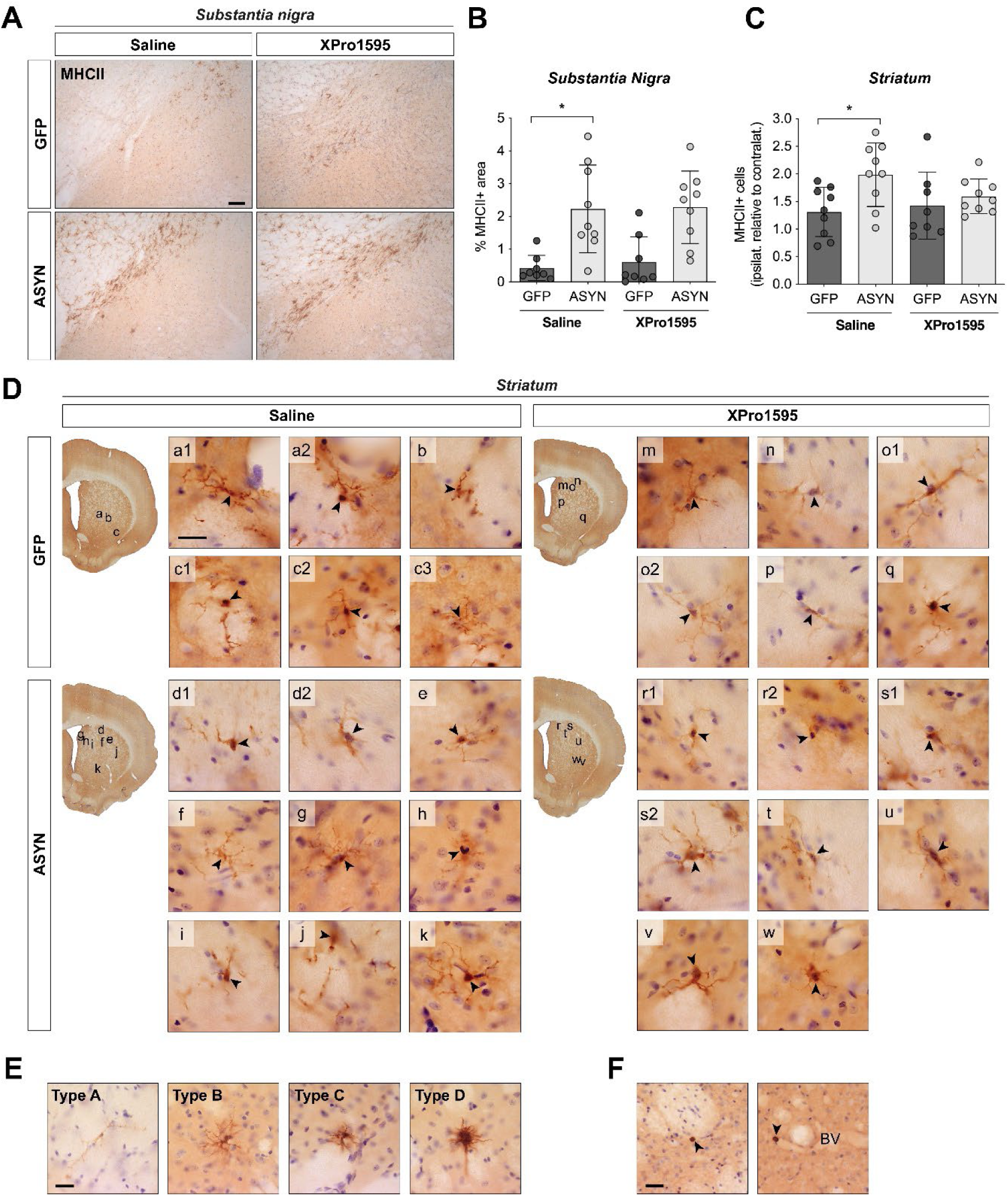
MHCII+ microglia in the Substantia nigra and Striatum. **A** Representative images of MHCII+ cells in the ipsilateral SN. **B** Bar graphs illustrates the percentage of area covered by MHCII+ immunostaining in the SN. **C** Bar graph showing the ipsilateral/contralateral ratio of striatal MHCII+ microglia numbers. Histogram presents the mean ± SD; two-way ANOVA followed by Tukey’s multiple comparisons test. * p<0.05 (*n* = 8-9/group). **D** Images showing MHCII+ microglia (arrowheads) in the ipsilateral side of a representative striatal section from each group. Lower case letters indicate the striatal localization of the cells. Scale bar: 20 µm (applies to all). **E** Example images showing the characteristic morphology of A-, B-, C-, and D-type microglia. Scale bar: 20 µm (applies to all). **F** Representation of round MHCII+ cells (arrowheads) observed in the parenchyma or associated to a blood vessel (BV). Scale bar: 30 µm.

For an indirect measurement of phagocytic activity in myeloid cells, we quantified the percentage of area covered by CD68+ staining in both striatum and SN **(FIG. 7A,B)**. While no differences were found among treatment groups in the striatum **(FIG. 7A,E)**, α-syn overexpression induced a significant increase in CD68 immunostaining in the SN **(FIG. 7B,F)**. Also here, XPro1595 treatment failed to modify the CD68 upregulation induced by α-syn expression.

**FIG. 7.**
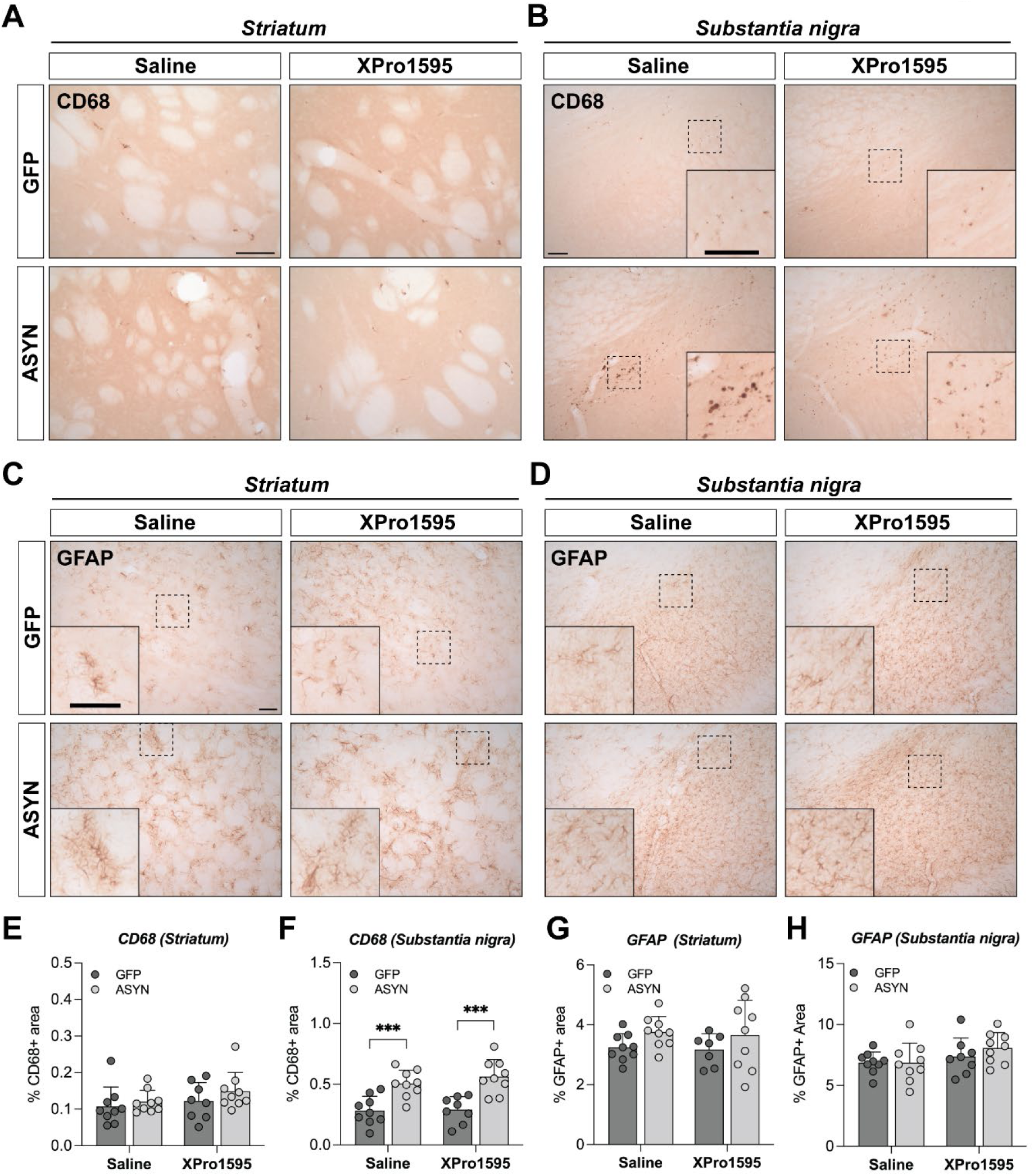
Immune response in the Striatum and Substantia nigra. **A-B** Representative images of CD68+ cells in the ipsilateral striatum and SN, respectively. **C-D** Representative images of GFAP+ cells in the ipsilateral striatum and SN, respectively. Scale bar represents 100 µm (applies to all). **E-F** Bar graphs showing the percentage of area covered by CD68+ immunostaining in the striatum (**E**) and SN (**F**). **G-H** Bar graphs showing the percentage of area covered by GFAP+ immunostaining in the striatum (**G**) and SN (**H**). The histograms present the mean ± SD (*n* =8-9/group). Two-way ANOVA followed by Sidak’s multiple comparisons test; *p<0.05, **p<0.01, ***p<0.001, ****p<0.0001.

Since astrocytes are also involved in the immune response in the brain, we evaluated if XPro1595 treatment would have any effect on the GFAP+ astrocytic response in the groups **(FIG. 7C,D)**. α-syn overexpression induced a small, non-significant increase in GFAP immunostaining in striatum with cells having a more activated morphological phenotype irrespective of the treatment **(FIG. 7C,G).** The GFAP expression in the SN was higher than that observed in striatum in all groups (see scale in **FIG. 7G,H**), but no significant differences between groups were observed **(FIG. 7D, H)**.

## DISCUSSION

TNF has been implicated as one of the cytokines mediating neuroinflammation while promoting neurodegeneration in PD (McCoy et al., 2008). Indeed, TNF levels are elevated in the brain, CSF and sera of PD patients (Eidson et al., 2017). Furthermore, circulating TNF levels have been positively correlated with the occurrence and severity of non-motor symptoms in PD patients (Lindqvist et al., 2012; Menza et al., 2010). Increased TNF levels in the nigrostriatal system have been reported in the 6-OHDA (Mogi et al., 1999) and MPTP toxin PD models, as well as in the rAAV-ASYN PD model (Chung et al., 2009; Theodore et al., 2008). In cellular and animal models, an excess of TNF leads to neuronal death (Neniskyte et al., 2014; van Loo and Bertrand, 2023). The use of DN-TNF biologics in PD is of ongoing interest, yet evidence regarding their effectiveness is still limited. Nevertheless, our previous studies showed the neuroprotective potential of the DN-TNF biologic XPro1595 in the toxic 6-OHDA PD model when administered intracerebrally (McCoy et al., 2006; McCoy et al., 2008), and peripherally (Barnum et al., 2014). Hence, in this study we aimed to evaluate the possible neuroprotective potential of peripheral XPro1595 treatment against α-syn-induced dopaminergic nigrostriatal neurodegeneration. To do so, we administered subcutaneous injections of XPro1595 (10 mg/kg) every 3 days for 12 weeks in the rAAV-ASYN PD model, starting 3 days post-rAAV injection. We assessed the motor behavior of the animals and evaluated the dopaminergic neuronal loss, α-syn pathology and immune response using histology. α-Syn overexpression did not result in overt motor defects in the model. α-Syn overexpressing rats showed pathological accumulation of phosphorylated α-syn in striatal axonal terminal swellings and SN neurons, dopaminergic axonal loss and SN VMAT2+ neuronal death, which were not prevented by the XPro1595 treatment. α-Syn pathology was paralleled by Iba1+ microgliosis and increased MHCII, and CD68 expression in the SN and striatum. While XPro1595 decreased the MCHII striatal upregulation, it did not prevent microgliosis, but in fact, promoted more hypertrophic and ameboid microglia in the SN. Therefore, peripheral administration of XPro1595 slightly modulated the brain immune response but this did not avoid the α-syn pathology, nor did it prevent the degeneration of SN dopaminergic neurons.

As before, α-syn overexpression led to swelling of axons where we found aggregated α-syn and accumulated TH and, to a lesser extent, VMAT2. α-syn expression in dopaminergic neurons resulted in a decrease of approximately -30% of TH axons in the striatum and a less pronounced decrease of VMAT2+ axons (−17%). This might be due to a TH downregulation by α-syn (Baptista et al., 2003; Yu et al., 2004), additionally suggested by observations in post-mortem brain tissue from PD patients where decreased TH and Nurr1—an activator of TH promoter activity—immunostaining intensity was seen in SN neurons containing α-syn+ inclusions, but not those without inclusions (Chu and Kordower, 2007; Chu et al., 2006). For this reason, VMAT2+ was used for stereological analysis in SN. As seen before in the model, α-syn overexpression resulted in dopaminergic VMAT+ neuronal loss (−30%) and this was not prevented by XPro1595 treatment. We also observed a loss of nigrostriatal neuronal cell bodies and terminals in both GFP control groups, indicating some degree of non-specific toxicity. GFP is the most common control protein but can cause damage in certain conditions. Indeed, Ulusoy et al. have reported dose-dependent toxicity following rAAV-mediated expression of GFP in the SN resulting in marked loss of TH+ and VMAT2+ neurons (Ulusoy et al., 2009). I*n vitro,* high titers of rAAV vectors of various serotypes including rAAV6 encoding GFP induced dose-dependent cytotoxicity and reduced cell viability in neurons and astroglia. This effect was in part related to vector itself but also to the GFP transgene since rAAV vectors containing a frameshift mutation in the GFP gene displayed reduced toxicity (Howard et al., 2008). The relatively high rAAV titer used in this study might have caused some undesired cytotoxicity. However, GFP groups showed no pSer129 α-syn pathology and no signs of axonal damage in the ipsilateral striatum despite the ∼19% loss of VMAT2+ SN neurons. This was not avoided, but rather non-significantly enhanced by peripheral administration of XPro1595, which might suggest that a certain level of inflammation might be beneficial to avoid this GFP toxicity.

In this study, α-syn overexpression induced mild motor changes in the stepping test but not in the cylinder test (despite some rats showing clear paw bias, see Fig. 1). We have previously seen motor asymmetry in both tests, using the rAAV2/5 α-syn that did not show anterograde spreading to striatum (Febbraro et al., 2013a; Febbraro et al., 2013b). The α-syn expression in striatal neurons seen herein might have induced neuronal dysfunction, complicating the interpretation of the motor behavior. Indeed, rAAV mediated A53T α-syn overexpression in ventral tegmental area (VTA) neurons resulted in the formation α-syn pathology and functional changes despite the lack of neuronal death (Maingay et al., 2006). *Snca*^+/+^ rat BAC full length human α-syn show loss of calbindin medium spiny neurons (MSN) and parvalbumin interneurons in striatum (Paldino et al., 2022). α-Syn pathology in striatal neurons has been previously described in the zQ175 Huntington’s disease transgenic line (Yu et al., 2022). Since the D2-MSNs are particularly sensitive to the accumulation of huntingtin, which are the first one to die in Huntington’s disease, one might speculate this could also be true for α-syn aggregation (Waldvogel et al., 2012). If so, α-syn-induced dysfunction of D2-MSNs could in theory cancel out the functional outcome of the reduced dopamine signaling and counteract motor impairments caused by the nigrostriatal degeneration. Hence, loss/dysfunction of MSNs could explain at least partially the lack of α-syn-induced motor impairment. On the other hand, GFP animals showed certain loss of SN neurons that could also justify the lack of robust differences between GFP and ASYN.

As before, we observed an α-syn-induced increase in SN microglia numbers, which show hypertrophic/ameboid morphology and upregulated CD68 and MHCII (Sanchez-Guajardo et al., 2010)). MHCII upregulation was also seen in the striatum. However, no changes in GFAP expression were seen in the model. Despite the lack of neuroprotection, the ASYN/XPro1595 group showed a differential immune response in the brain, with more hypertrophic Iba1+ cells in SN (increase in C and D type) and a trend towards less MHCII expression in the striatum. This suggests that although the XPro1595 treatment influenced the inflammatory reaction, this was not protective. It remains unclear whether this effect is exerted directly in the brain or in peripheral immune cells that in turn influence the brain upon infiltration. Indeed, the hypertrophic/ameboid Iba1+ cells analyzed are indistinguishable from infiltrated macrophages. Although no macroscopic changes were observed in the spleen, we cannot discount changes in peripheral immune cell functionality.

Since neuroprotection was not achieved in the present study, it is relevant to address the administration route. In this study, XPro1595 was administered subcutaneously following the dosing regimen applied by us previously in rats of the same strain and age, but opposite sex (Barnum et al., 2014). There, XPro1595 was detected in serum (4787±1931 ng/ml) and CSF (3±1 ng/ml) using an anti-human TNF immunoassay, thus we anticipate similar levels in the present study. Two studies have shown protective effects of systemic XPro1595 treatment in the experimental autoimmune encephalomyelitis model of multiple sclerosis (Brambilla et al., 2011; Taoufik et al., 2011). Another study has shown an early boost of remyelination and improved phagocytosis of myelin debris by CNS macrophages in the cuprizone model of multiple sclerosis, despite not preventing toxin-induced oligodendrocyte loss and demyelination (Karamita et al., 2017). In mild-to-moderate traumatic brain injury (CCI mouse model), systemic XPro1595 treatment improved hippocampal pathology and neurological recovery (Larson et al., 2022). In an Alzheimer’s disease mouse model, peripheral XPro1595 administration decreased overall CD4+ T cell numbers and activated myeloid-derived immune cells (MacPherson et al., 2017). All these studies employed a similar dosing regimen with subcutaneous injections of XPro1595 (10 mg/kg) every third day or twice weekly, resulting in decreased neuropathology and reduced levels of pro-inflammatory cytokines. Moreover, a single intravenous injection of XPro1595 was found to induce partial protection against focal cerebral ischemia; the treatment failed to reduce infarct size, but did improve motor performance correlating with decreased infiltration of peripheral immune cells (Clausen et al., 2014). However, there are contrasting studies where protective effects were achieved with central, but not systemic XPro1595 treatment or vice-versa. In a mouse model of spinal cord injury, epidural infusion of XPro1595 significantly reduced the lesion size and improved the clinical outcomes; however, these effects were not achieved via subcutaneous injection of XPro1595 (10 mg/kg) every third day (Novrup et al., 2014). Yet, another more recent study has shown that continuously epidurally delivered XPro1595 (micro-osmotic pumps) decreased IL-1β and IL-6 levels and increased IL-10 in the acute phase after spinal cord injury, followed by reduction of infiltrated macrophages and neutrophils and modulation of microglia activation, improving functional outcomes in the model (Lund et al., 2023). Likewise, in a model of Huntington’s disease, chronic infusion of XPro1595 (0.08 mg/kg/day) into the lateral ventricle significantly reduced gliosis, rescued neuronal density, reduced huntingtin aggregation, and improved motor function; however, intraperitoneal injection of XPro1595 (30 mg/kg) twice weekly failed to induce protection probably due to systemic degradation and insufficient concentrations of XPro1595 reaching the brain (Hsiao et al., 2014). In the study, following systemic administration, XPro1595 was detected in the serum and in the liver but not in the cortex or striatum. Conversely, after central administration, the drug was detected in the cortex and striatum, but not in serum or liver. Central administration significantly reduced TNF levels and caspase activation in the brain, while these measures were not affected by systemic drug administration (Hsiao et al., 2014). The importance of achieving a sufficient drug concentration in the affected brain area was corroborated by us in a study showing that infusion of the DN-TNF drug XENP345 (0.08 mg/kg/day) into the SN of the 6-OHDA model resulted in marked rescue of SN dopaminergic neurons and attenuation of motor behavior. Strikingly, no protective effects were observed, when the drug was infused into the striatum, indicating that a high drug concentration locally in the SN is crucial for achieving protection in this PD model (McCoy et al., 2006). In another study we have highlighted that XPro1595 exerts significant peripheral effects which may not translate effectively to central neuroprotection. In the MPTP non-human primate model of PD, despite the (sexually dimorphic) responses in microglia activation ([^18^F]FEPPA PET scans to evaluate translocator proteins) and motor deficits, solTNF inhibition led to attenuated inflammation in biofluids and reduced CD68 expression in the lower colon, but was not able to protect against escalating doses of MPTP-induced neuronal loss (Joers et al., 2020). In humans with mild cognitive impairment enrolled in a Phas1b clinical trial, XPro1595 was reported to improve synaptic health markers (see www.inmunebio.com).

One limitation of our study is the exclusive use of female rats, which may have influenced the outcome compared to other studies with mixed or male cohorts. The impact of sex differences on the response to TNF inhibition has been suggested but not extensively explored. As mentioned, in the MPTP non-human primate model of PD, sex-dimorphic effects were observed, with males exhibiting more severe parkinsonism than females and differing microbiota diversity, which could confound the evaluation of responses to TNF inhibition (Joers et al., 2020). We and others have reported sex-specific differences in α-syn toxicity and immune response (Ferreira et al., 2023; Lamontagne-Proulx et al., 2023), highlighting the need to consider these variations in future research. Despite the lack of strong motor defects, however we successfully modeled the two hallmarks of PD α-syn aggregation and SN degeneration, thus we believe our study holds significant relevance for the PD field. In conclusion, our study shows that peripheral administration of the DN-TNF XPro1595 at doses that have been previously shown to confer CNS neuroprotection in other neurodegenerative models was not protective in the AAV-α-syn PD model. This might be due to the requirement for higher drug levels to achieve protection in this specific model. Alternatively, the inhibition of TNF signaling might be protective only if targeted during specific stages of the neurodegenerative process. Understanding the temporal interplay between neurodegeneration and neuroinflammation may elucidate why neuroprotection was not observed in this model, in contrast to the 6-OHDA model.

## CONTRIBUTIONS

Conceptualization; MRR & MGT. Formal analysis and investigation; JRC, SAF & MRR. Resources; JJ & DES. Funding acquisition, project administration and supervision; MRR. Writing - original draft; JRC, SAF & MRR. Writing - review & editing; all authors

## Supplementary figures

**FIG. S1.**
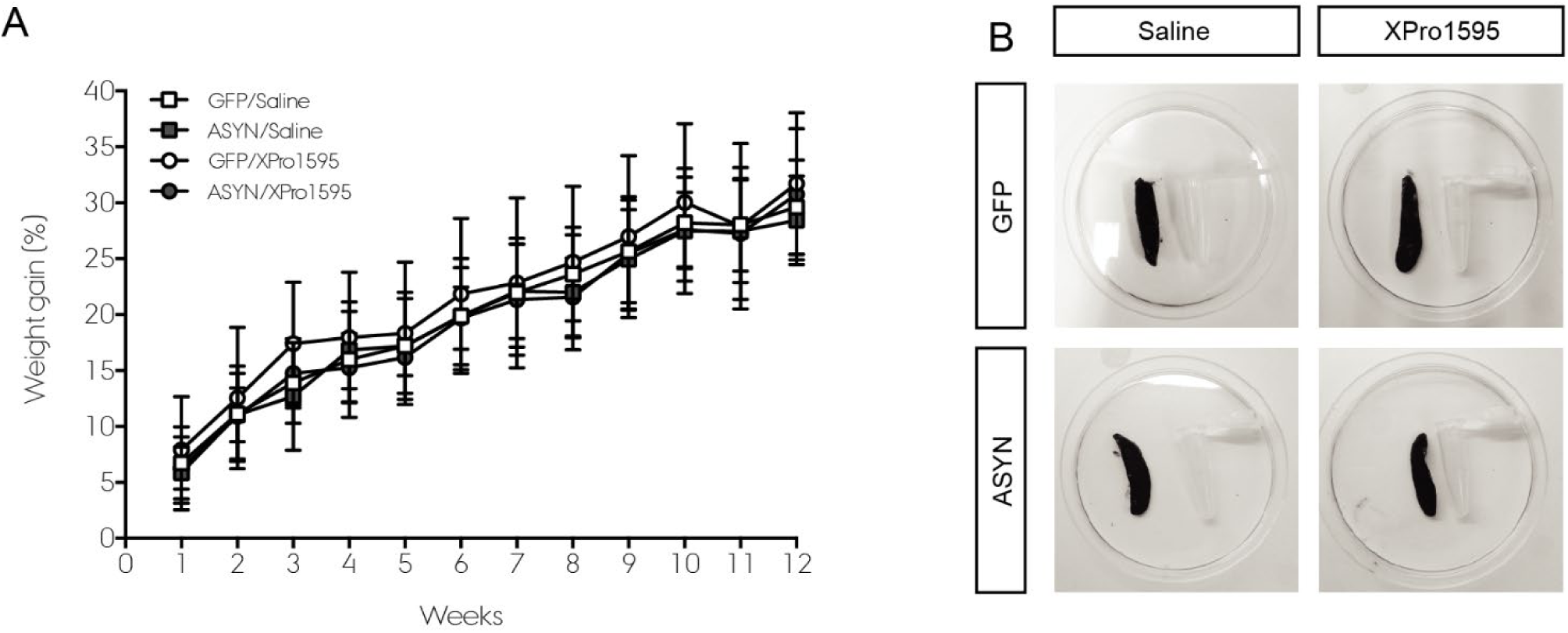
Systemic effects. Neither body weight nor spleen size were affected by the rAAV-injection and pharmacological treatment. **A** Percentage of weight gain during the 12-week study period. **B** Representative photos of a spleen from each experimental group. An eppendorf tube was included as an indicator of size.

**FIG. S2.**
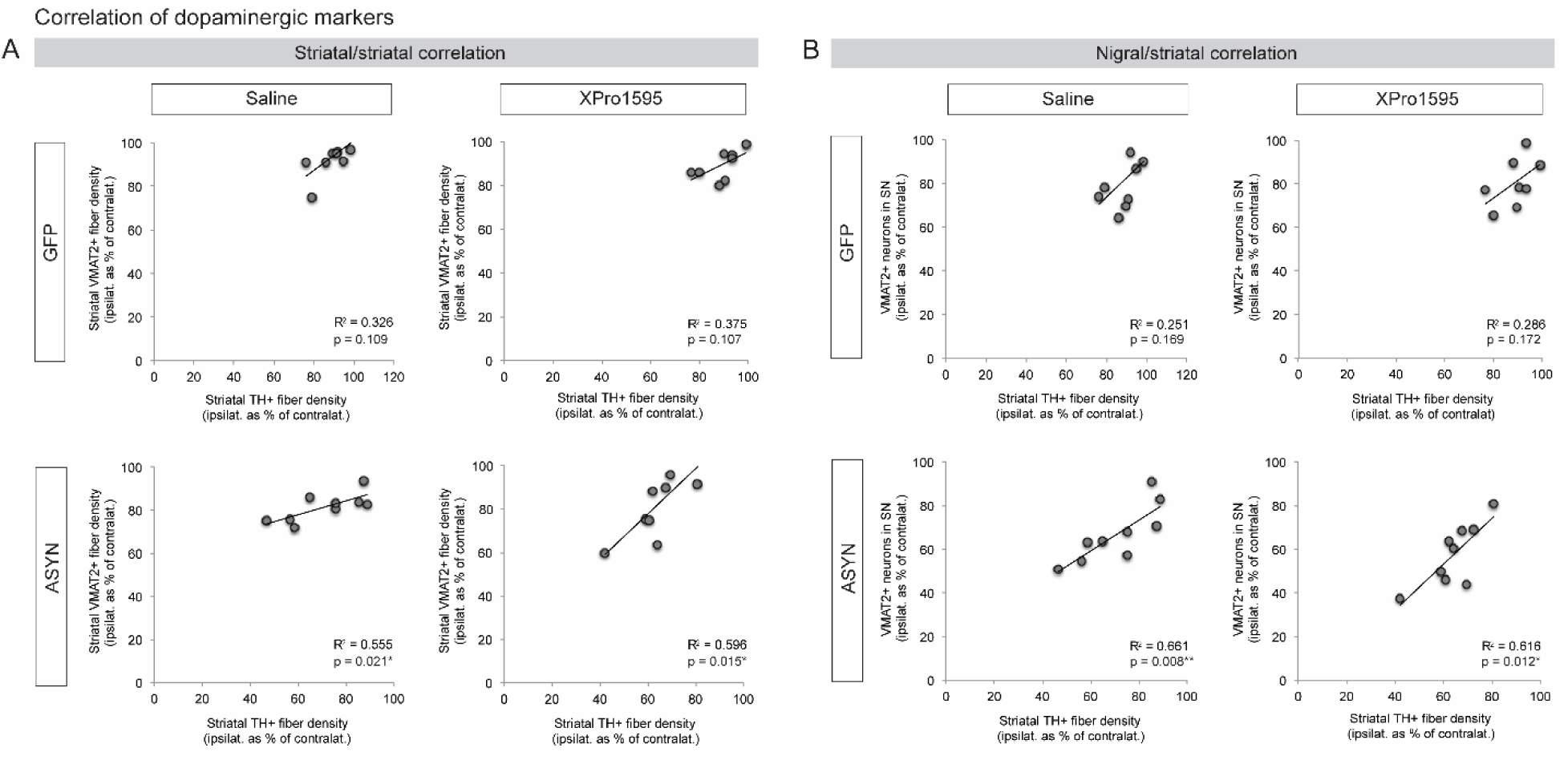
Correlation of dopaminergic markers. **A** Correlation of the striatal TH+ and VMAT2+ fiber densities. **B** Correlation of the striatal TH+ fiber density and the number of VMAT2+ neurons in SN.

